# HapCNV: A Comprehensive Framework for CNV Detection in Low-input DNA Sequencing Data

**DOI:** 10.1101/2024.12.19.629494

**Authors:** Xuanxuan Yu, Fei Qin, Shiwei Liu, Noah J. Brown, Qing Lu, Guoshuai Cai, Jennifer L. Guler, Feifei Xiao

**Affiliations:** Department of Epidemiology and Biostatistics, Arnold School of Public Health, University of South Carolina, Columbia, SC, USA; Division of Cancer Epidemiology and Genetics, National Cancer Institute, 9609 Medical Center Drive, Rockville, MD, 20850, USA; Center for Neuroimaging, Department of Radiology and Imaging Sciences, Indiana University School of Medicine, Indianapolis, Indiana, USA; Department of Biology, University of Virginia, Charlottesville, VA, USA; Department of Biostatistics, College of Public Health and Health Promotions & College of Medicine, University of Florida, Gainesville, FL, USA; Department of Surgery, College of Medicine, University of Florida, Gainesville, FL, USA

**Keywords:** Single-cell DNA sequencing, Low-input sequencing, Copy number variation, Haploid, Pseudo-reference sequence

## Abstract

Copy number variants (CNVs) are prevalent in both diploid and haploid genomes, with the latter containing a single copy of each gene. Studying CNVs in genomes from single or few cells is significantly advancing our knowledge in human disorders and disease susceptibility. Low-input including low-cell and single-cell sequencing data for haploid and diploid organisms generally displays shallow and highly non-uniform read counts resulting from the whole genome amplification steps that introduce amplification biases. In addition, haploid organisms typically possess relatively short genomes and require a higher degree of DNA amplification compared to diploid organisms. However, most CNV detection methods are specifically developed for diploid genomes without specific consideration of effects on haploid genomes. Challenges also reside in reference samples or normal controls which are used to provide baseline signals for defining copy number losses or gains. In traditional methods, references are usually pre-specified from cells that are assumed to be normal or disease-free. However, the use of pre-defined reference cells can bias results if common CNVs are present. Here, we present the development of a comprehensive statistical framework for data normalization and CNV detection in haploid single- or low-cell DNA sequencing data called HapCNV. The prominent advancement is the construction of a novel genomic location specific pseudo-reference that selects unbiased references using a preliminary cell clustering method. This approach effectively preserves common CNVs. Using simulations, we demonstrated that HapCNV outperformed existing methods by generating more accurate CNV detection, especially for short CNVs. Superior performance of HapCNV was also validated in detecting known CNVs in a real *P. falciparum* parasite dataset. In conclusion, HapCNV provides a novel and useful approach for CNV detection in haploid low-input sequencing datasets, with easy applicability to diploids.

## Introduction

Variations in the copy number of consecutive genomic regions, termed copy number variants (CNVs), are important adaptive strategies and a major source of phenotype diversity in many organisms [1–5]. CNVs have been appreciated as drivers for many human complex disorders such as lung cancer [6, 7] and breast cancer [8–10]. CNVs are also important in driving evolution of microbes such as *Escherichia coli* [11, 12] and *S. cerevisiae* [13–15]. In *S. cerevisiae*, duplications of the *GAP1* gene are repeatedly generated in different nitrogen-limited chemostat environments [16]. The haploid protozoan parasite *Plasmodium falciparum*, which is the most fatal human malaria parasite and contributes to a large number of deaths annually [17, 18], readily evolves CNVs in response to antimalarial drug pressure [19]. Consequently, specific CNVs such as the multi-drug resistance-1 gene, *pfmdr1*, attain high frequencies within parasite populations [20–24].

Novel CNVs arising in a predecessor genome offer insights into the evolutional potential of genome under stress or selection. These rare, or *de novo,* CNVs are often deleterious and exist in small populations [25], and their rates may increase under stress, which was evidenced by the studies of adaptive evolution of microbes [26, 27]. In humans, rare CNVs are shown to be associated with increased risks of ovarian cancer and ischemic heart disease [28]. As such, detecting both common and rare CNVs will pave the way to answer several pertinent questions such as the evolution and adaption of these variants under stress in different species.

Low-input DNA sequencing including low-cell and single-cell sequencing (scDNA-seq) has enabled the characterization of genomes at a cellular level, providing an unprecedented opportunity to detect and understand the roles of CNVs in diverse phenotypes and population structure [29]. Challenges of the discovery of CNVs arise in two aspects. First, the procedure of whole genome amplification introduces amplification biases and dropout events, resulting in extremely shallow and highly non-uniform read counts [30, 31], especially when the input DNA amount is low. This presents a substantial challenge for accurate CNV detection in haploid organisms because of the relatively short genome and a higher level of DNA amplification [32]. Second, how to select suitable normal controls for normalizing read-depth data is a historical challenge. In traditional CNV detection methods, reference cells are usually pre-specified from cells that are assumed to be normal or disease-free. For example, SCOPE [30] and SCNV [33] identify the normal control or constructs composite reference cells from a set of cells with low signal variations. Utilizing these strategies, signals from common CNVs shared among reference and test cells are canceled out, leading to inflated false negatives during CNV detection. False positives may be similarly generated if the reference contains rare CNVs that are absent from the majority of cells. When “normal cells” are unavailable, attempts have been made to construct pseudo-reference cells [34, 35]. However, they either use within-tested cell information or average information from all cells, which introduces bias because the reference cells are still mixture of different copy number states. Due to these limitations, common CNVs (population frequency > 10%) are largely overlooked even though detection of rare CNVs (population frequency < 10%) is less susceptible.

To overcome these limitations and perform globally accurate CNV detection, we developed a comprehensive statistical framework for CNV detection with low-input haploid data called HapCNV. We introduced a novel genomic location-specific pseudo-reference construction strategy that selects unbiased references using a preliminary cell clustering method. This cell clustering method uses the copy number profile to define cells that are in a “normal state” (i.e., no copy number gain or loss) for each genomic location. Utilizing this strategy for data normalization, we systematically alleviated amplification biases and effectively retained both common and rare CNVs. We comprehensively evaluated our method in simulated and a real dataset and verified its superior performance by benchmarking to existing methods. Our results demonstrated the exceptional performance of our method in detecting both common and rare CNVs.

## Methods

### Data and notations

Our method models the read count matrix that contains *M* genomic bins or markers and *N* cells. Binning is a pre-processing step where the read counts of data are summarized in a large unit instead of per base pair, whereas the length of each bin is defined by accommodating the needs of each study. In our applications, we used 1kb (i.e., 1000 base pairs) as the bin size in the binning process. It is found that there is a large abundance of small CNVs (<300bp) in the haploid genome [36, 37], users can accommodate to the study and define small windows in the binning process. 𝑥_𝑖𝑗_ denotes the number of read counts for the 𝑖-th (𝑖 = 1, … , 𝑀) bin from the 𝑗-th (𝑗 = 1, … , 𝑁) cell.

### Sample quality control and bias correction

First, we removed genetic markers or bins of poor quality (calling rate < 80%) or bins with alleviated bias arising from technical artifacts and biases introduced in the sequencing process (i.e., GC content less than 20% or over 80%). Bins with ambiguous or potentially incorrect mapping of reads were removed for those with mappability < 0.9 (Fig. 1A, 1B).

**Fig. 1.**
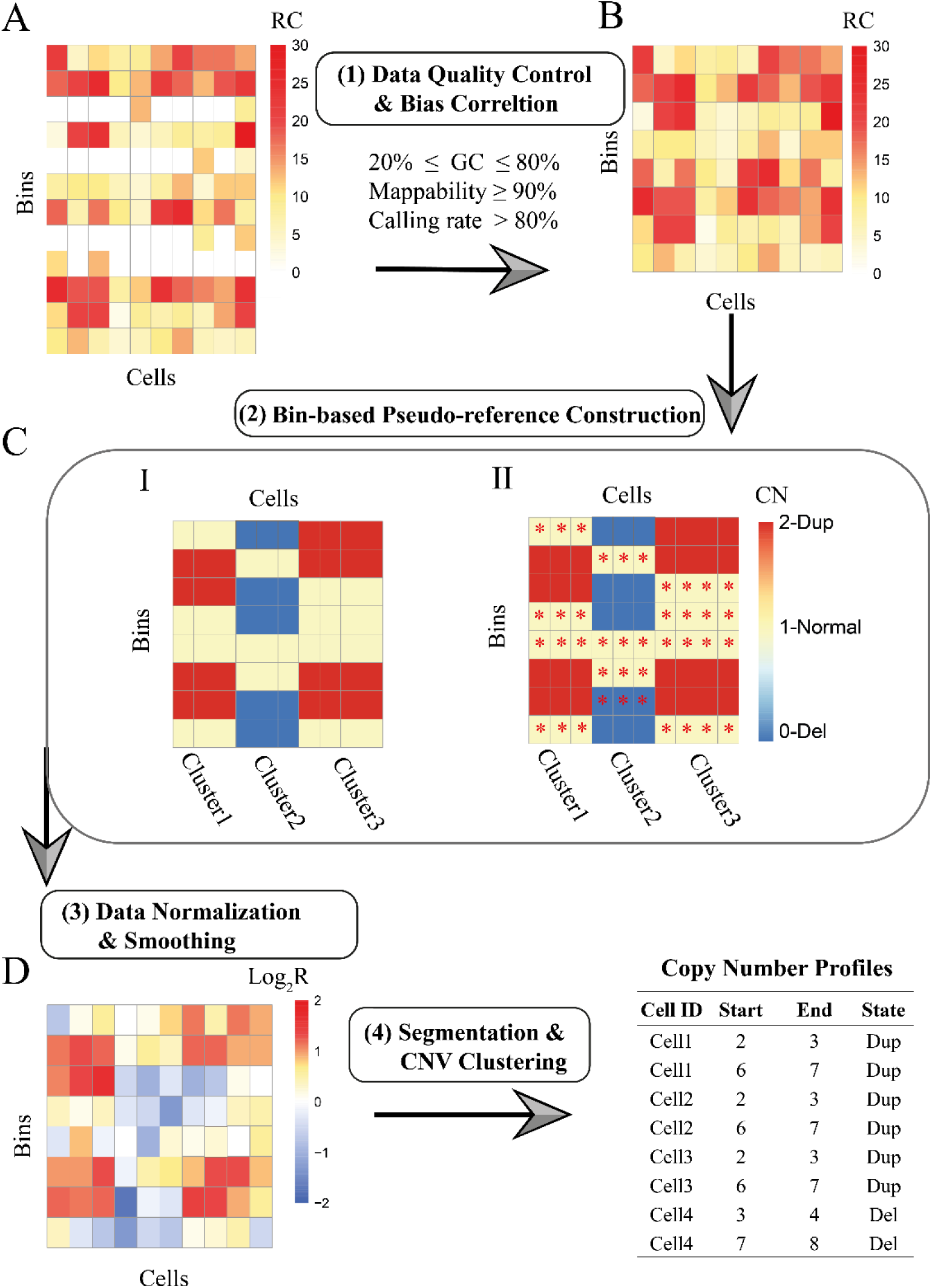
Flowchart of HapCNV. **A**: Low-cell or single-cell DNA-seq raw data matrix with rows representing genomic markers or bins, and columns denoting cells. **B**: Read count matrix after quality control and correction of biases introduced by GC content and mappability. **C**: Pseudo-reference construction: the left panel shows the results of cell clustering, where three clusters are identified with shared CNVs detected within clusters; the right panel shows the reference sequence constructed by utilizing information from cells marked with red stars. For example, when constructing reference sequence for the first cell per bin, other cells identified to be normal states will be used as the references. Specifically, for the first bin, the two star-labeled normal cells will serve as the reference, the median of which will be used to normalize the read count of the first cell. We considered three copy number states: 2-duplications; 1-normal states; 0-deletions. **D**: The logarithm transformation is conducted after the normalized matrix is calculated by dividing the read counts of each tested cell over the median of the read counts from the pseudo-reference sequences. An additional smoothing step is further conducted to remove outliers in the intensity data. **E**: Segmentation and CNV clustering are conducted using Circular Binary Segmentation (CBS) method and Gaussian Mixture Model (GMM), separately.

We then conducted a two-step normalization procedure to further remove the bias associated with mappability and GC content, that indicate the sequencing efficiency and alignment accuracy of genome regions. Such strategy has been utilized in existing diploid CNV detection methods [38, 39].

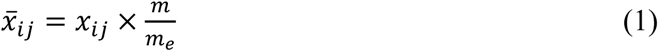

where 𝑚_𝑒_ is the median read count of all the bins sharing the same 𝑒 value (𝑒 = GC content and mappability) as the 𝑖-th bin, and 𝑚 is the global median read count of all the bins. The library size was also adjusted to ensure that all cells have equivalent total read counts, effectively mitigating differences in the sequencing depths.

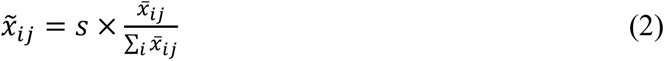

where 𝑠 is the scaling factor for cell level normalization, such as 1,000,000.

### Segmentation and classification of copy number profiles

After pseudo-reference construction based on shared CNV profiles and cell clusters generated using FLCNA [38], logarithm transformation was conducted by dividing 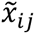 of the tested cell to the median of the pseudo-reference read counts for each bin. To efficiently reduce the noise resulting from outliers, a sequence smoothing method [40] was systematically applied with the normalized data where the outlier along the sequence was replaced by the median of those from the neighboring bins. Specifically, with a window size of *B* and a threshold of *t*, the normalized signal 𝐿𝑜𝑔_2_𝑅 was considered an outlier if it deviated from the window mean by more than *t* times of the within-window variation. It has been shown that CBS algorithm is an efficient approach for changed point detection [41–43] which has been widely applied in numerous studies [44–47]. We utilized the “DNAcopy” package [48] to implement the CBS method for detection of change points in each cell. Ultimately, a Gaussian Mixture model (GMM) [40] was utilized for CNV states identification which was extensively utilized in our previous studies [38, 40, 49] to assign the most likely CNV state to each segment. Given the variability of mean signals for CNV states across species or datasets, it is crucial to achieve accurate assignment of initial signal means to maximize the precision of copy number state assignment. In this case, we generated the data-specific initial means by clustering the segmental means estimated by CBS using “mclust” (version 6.1.1) R package [50] and then used GMM to update the segment means for each copy number state.

### Evaluation of HapCNV via simulations

To validate the theoretical advantage of HapCNV in efficient CNV detection globally, we commenced our assessment for detection of a wide range of CNV frequencies in the population including both common (> 10% frequency in the population) and rare CNVs (<10% frequency in the population). We conducted *in-silico* simulation studies benchmarking to existing methods including SCOPE, HMMcopy, and SCCNV [30, 34, 35]. DNAcopy [41] was not included in the list of tools for comparison because it is not applicable to the simulated read count data without specification of reference cells. The effect of varied CNV lengths (short: 3∼10 bins, medium: 10∼50 bins, and long: 50∼100 bins), copy numbers, proportions of cells with the variants, and combinations of different copy number states on the performance of these methods was evaluated. Precision rate 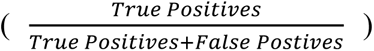 recall rate 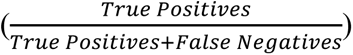, and F1 score 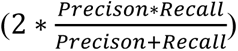 were used for the assessment.

For CNV data generation mimicking real data, we first generated CNV-free data using a scDNA-seq dataset comprised of genomes from 28 haploid early erythrocytic stage *Dd2 P. falciparum* parasites [19]. In this previous study, individual parasite genomes were amplified using an optimized MALBAC protocol and sequenced using the Illumina Nextseq 550 (150bp paired-end cycles). After genomic binning and quality control filters based on GC content, mappability, and calling rate, we obtained 15,718 genomic bins (defined in 1,000 bps) or markers across 14 chromosomes. HMMcopy [34], Lumpy [51], and Ginkgo [52] were used to initially identify regions with CNVs. These specific regions were then substituted with data mimicking normal copy number states.

Note that amplification bias was introduced during whole genome amplification, leading to bin-specific biases in read depth. To remove the potential effect of amplification bias on the performance of our methods, we also generated CNV-free data by randomly shuffling the CNV-free background sequences across genomic positions. Due to the small sample size in the actual dataset, we augmented the background sequences four times to increase the sample size.

With the generated CNV-free background sequences (either unshuffled or shuffled), we randomly chose locations of 80 CNV regions (CNVRs) across the genome with varied CNV lengths (short: 3∼10 genomic bins, medium: 10∼50 bins and long: 50∼100 bins). Since HapCNV involves initial identification of cell clusters for pseudo-reference construction, the cells were randomly assigned to two clusters with each cluster characterized by 40 CNVRs. For each CNVR, cells were randomly assigned to contain CNVs with varied proportions in the simulated sample (i.e., 5%∼90%). Finally, CNV signals were spiked in by multiplying the background signal by *c*/2, where *c* was a normal random variable generated from 𝑁(0.4, 0.1^2^) for deletion of a single copy (DEL), or 𝑁(4.2, 0.1^2^) for duplication of a single copy (DUP) [30]. We also included a scenario involving a mixture of different copy number states (MIXED) with different signal strength (i.e., DEL and DUP). After the logarithm transformation, the CNVs with DEL showed stronger signal than DUP (mean of −2.32 versus 1.07).

### Application of HapCNV to low-cell sequencing data of *P. falciparum* parasite

We used an independent low-cell DNA sequencing dataset from haploid early erythrocytic stage *P. falciparum* parasites that consisted of 39 samples and 23, 299 genomic bins (a bin defined in 1,000 bps) per genome [32]. Due to the small, AT-rich genome (23 Mb, 19.4% GC [53]) of *P. falciparum* malaria parasites, single-cell genomics is challenging [19]. In this more recent study, small numbers of parasites from two parasite lines were isolated for low-cell genomics (i.e., two cells were sequenced together in each sequencer well). Genomes were amplified using a further optimized MALBAC protocol and once again sequenced using the Illumina Nextseq 550 (150bp paired-end cycles) [32]. We used data from 27 untreated *Dd2* samples and 12 *FCR3* samples for our analysis.

We used *Dd2* samples to evaluate various CNV calling methods (i.e., HapCNV, DNAcopy, SCOPE, HMMcopy, and SCCNV) in detecting the frequencies of rare (≤ 10%), common (> 10%), and two known CNVs (*pfmdr1* CNV: chr5: 888001-970000, ∼82kb and *pf332* CNV: chr11: 1950201-1962400, ∼12kb) [19, 32]. Since the *pfmdr1* CNV was not detected in the *FCR3* cell line, we used *FCR3* samples as references for DNAcopy and SCOPE, which require manual selection of control samples to identify known and global CNVs (including common and rare ones).

Data preprocessing and quality control were conducted in a similar manner across all methods. Default parameters were used in HMMcopy and DNAcopy. In SCCNV, we used probability thresholds ranging from 0.2 to 0.5 to define the CNV states to improve the sensitivity. For SCOPE, the minimal ploidy was set at one to accommodate haploid data, and one latent factor was included in the normalization to capture potential biases.

The CNVs identified from different methods had inconsistent genomic boundaries that made comparisons difficult. To further refine CNVs and classify them as common or rare, we summarized the identified variants using the CNVR defining algorithm implemented in the CNVRuler software [54]. As such, the data-driven CNVRs were constructed that use the same genomic boundaries among samples, consequently, each sample presented duplications, deletions, or haploid. Based on the defined CNVRs, precision rate, recall rate, and F1 score were computed for assessment of the methods.

## Results

### Summary of HapCNV workflow

The HapCNV framework aimed to detect CNVs in haploid single-cell or low-cell genomes. It consisted of four major steps (Fig. 1): (i) data pre-processing including quality control and bias correction; (ii) bin based pseudo-reference construction; (iii) data normalization and data smoothing to remove outliers; and (iv) variant calling including genomic segmentation for breakpoints identification and copy number state estimation. In summary, starting with raw data, we first retained genomic bins with good data quality or those with less bias due to technical factors. To construct the pseudo-reference for normalization, we used our previously developed single-cell clustering tool, FLCNA [38], to cluster cells and identify CNVs shared within cluster. For each bin, we identified clusters with normal states (i.e., diploids with no duplications or deletions); then cells within the cluster served as the reference for all other cells under investigation. These genome-wide bin-specific references were called pseudo-reference. We used these references to construct logarithm-transformed intensity data, followed by chromosomal segmentation and classification of copy number profiles (details in methods).

For the computational speed of HapCNV, using a high-performance computing cluster with a 32-Core Processor and 540 GB of RAM, it took about 40 minutes to complete the analysis of a dataset with 69 haploid cells.

### Bin-based pseudo-reference construction

For CNV detection, reference cells or normal controls are desirable to provide a baseline to define copy number gain or loss of the genomic segments. Properly constructed references aid in the removal of amplification biases while preserving CNV signals [35]. However, using inappropriate references may lead to signal cancellation or reversion of the copy number states (Supplementary Fig.S1). Thus, existing CNV detection methods [30, 33, 35, 41] that rely on arbitrarily selecting cells or pre-defined normal controls may lose efficacy in detecting common CNVs. Herein we developed a strategy of bin-based pseudo reference construction to adaptively construct a sequence that will be used for data normalization (Fig. 1C). To do so, we used a method that we have previously developed for copy number profile-based single-cell clustering, FLCNA [38], to identify cells with normal states for each genomic bin. We then used cells that were identified to be normal states as reference cells for each genomic bin. For each test cell, such pseudo-reference was constructed for the whole genome and used as a baseline for test cell bins comparison. A special case was when there was no heterogeneous copy number state across all cells for a specific bin, the pseudo-reference was constructed using bins of normal state within the tested cell itself.

### Simulations showed superior performance of HapCNV in data without amplification biases

For the simulated data mimicking real data, when the original genomic structure from the cells was retained, amplification bias was evident. To remove this effect, we randomly shuffled the sequences before spike-in of CNV signals. With such data, HapCNV demonstrated superior performance in detecting CNVs of different sizes (short: 3∼10 genomic bins, medium: 10∼50 bins and long: 50∼100 bins) and copy number states (DEL, DUP, and MIXED) (Fig. 2). HapCNV showed strong overall performance for short CNVs with highest F1 scores (Fig.2A-C), highlighting the benefit of the pseudo-reference construction procedure using the pre-clustering results of the cells and preservation of signals from both rare and common CNVs. In contrast, the other methods had many false positive calls, leading to low precision rates and F1 scores (Fig. 2A-C, Fig. S3). For medium and long CNVs (Fig. 2D-I), HapCNV maintained the power in detecting CNVs in most scenarios. HMMcopy also showed a good performance in detecting long DEL and MIXED CNVs. However, without borrowing information across cells, HMMcopy showed slightly reduced performance as CNV proportion increased, which was especially evident when CNVs were shared by a large percentage of the cells. Collectively, by adaptively constructing pseudo-reference sequences using signals across cells, HapCNV consistently demonstrated superior performance over existing methods in detecting both rare and common CNVs, especially for short CNVs.

**Fig. 2.**
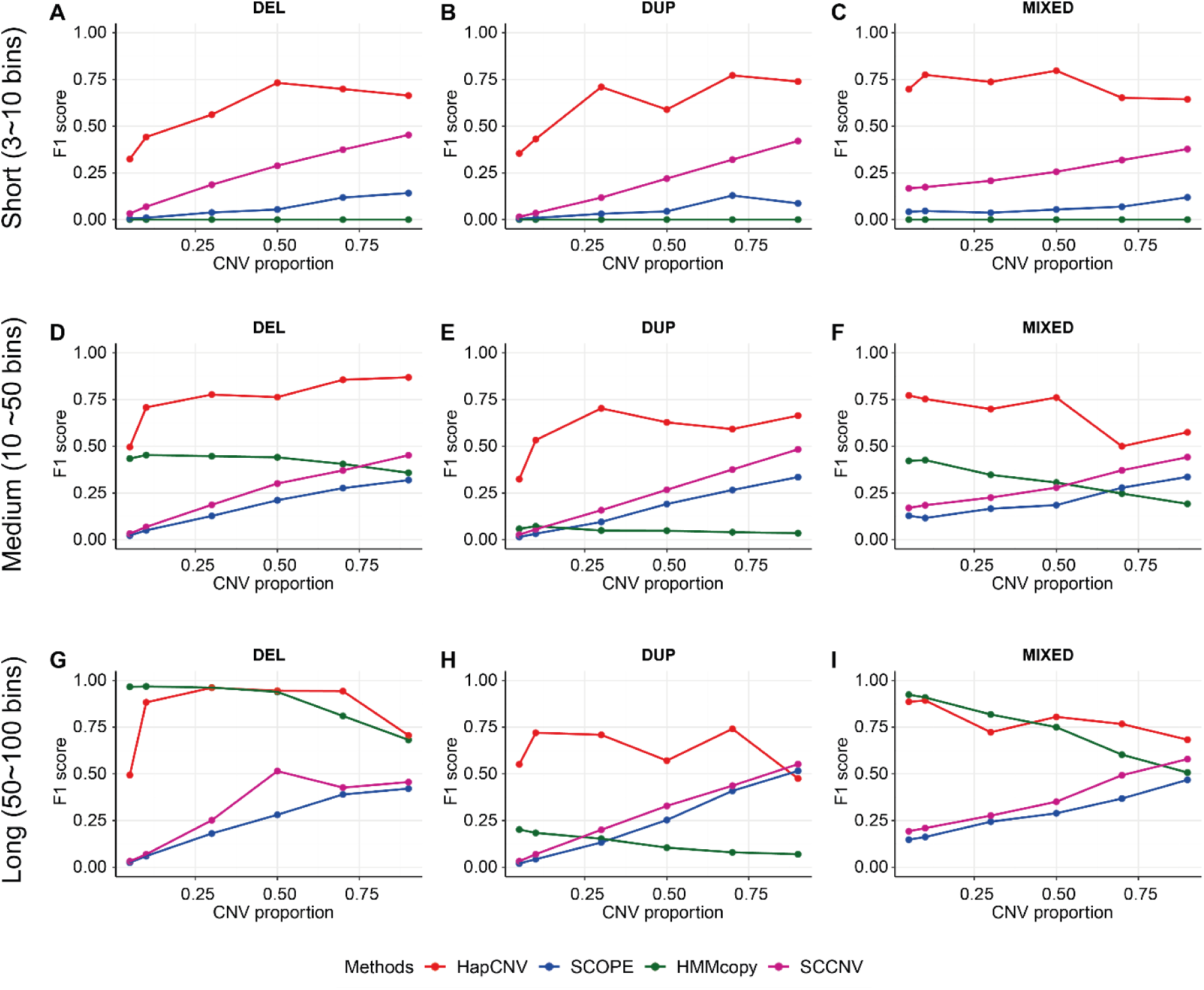
Performance evaluation of HapCNV using F1 score in comparison with existing methods for simulated data without amplification biases. For the simulations, we randomly shuffled the locations of the genomic biomarkers prior to spiking in the CNV signals to remove the effect that amplification biases may bring to the evaluation. Signals for 118 cells were simulated for three different CNV states: deletion of a single copy (DEL), duplication of a single copy (DUP), and mixture of two CNV states (MIXED) with varied length. The CNV proportion ranges from 5% to 90%.

### Simulations showed superior performance of HapCNV in data with amplification biases

Next, we evaluated the performance of HapCNV using simulations with amplification biases which retained the genomic locations of real data prior to spike-in of CNV signals. As expected, the performance of HapCNV was slightly impacted (Fig. 3) compared to the more ideal scenario when background noise was randomly shuffled across the genome (Fig. 2). Still, HapCNV demonstrated superior performance in detecting short CNVs (Fig. 3A-C), while retaining its ability to detect common CNVs for medium and long CNVs (Fig. 3D-I). Consistently, when CNV length was short or signals were weak (i.e., for duplications of a single copy [DUP]), HapCNV showed the best performance among these methods. This advantage became more evident as CNV proportion increased, underscoring the improved efficacy of HapCNV by the cell pre-clustering step for pseudo-reference construction. SCOPE and SCCNV still suffered from false positives (Fig. S4, Fig. S5). In contrast, although HapCNV and HMMcopy detected fewer true CNVs, they showed high precision rates (Fig. S4, Fig. S5). In detecting medium and long CNVs, HapCNV and HMMcopy demonstrated superior performance for CNVs of strong signals with higher precision rates than the other methods (Fig. 3, Fig. S4, Fig. S5). HMMcopy was found to be less sensitive to amplification biases and showed overall good performance for detecting medium or long CNVs with strong signals (Fig. 3D-I). Taken together, HapCNV presented the desired and stable performance in CNV detection, especially in detecting short CNVs and CNVs with weak signal when amplification biases were present.

**Fig. 3.**
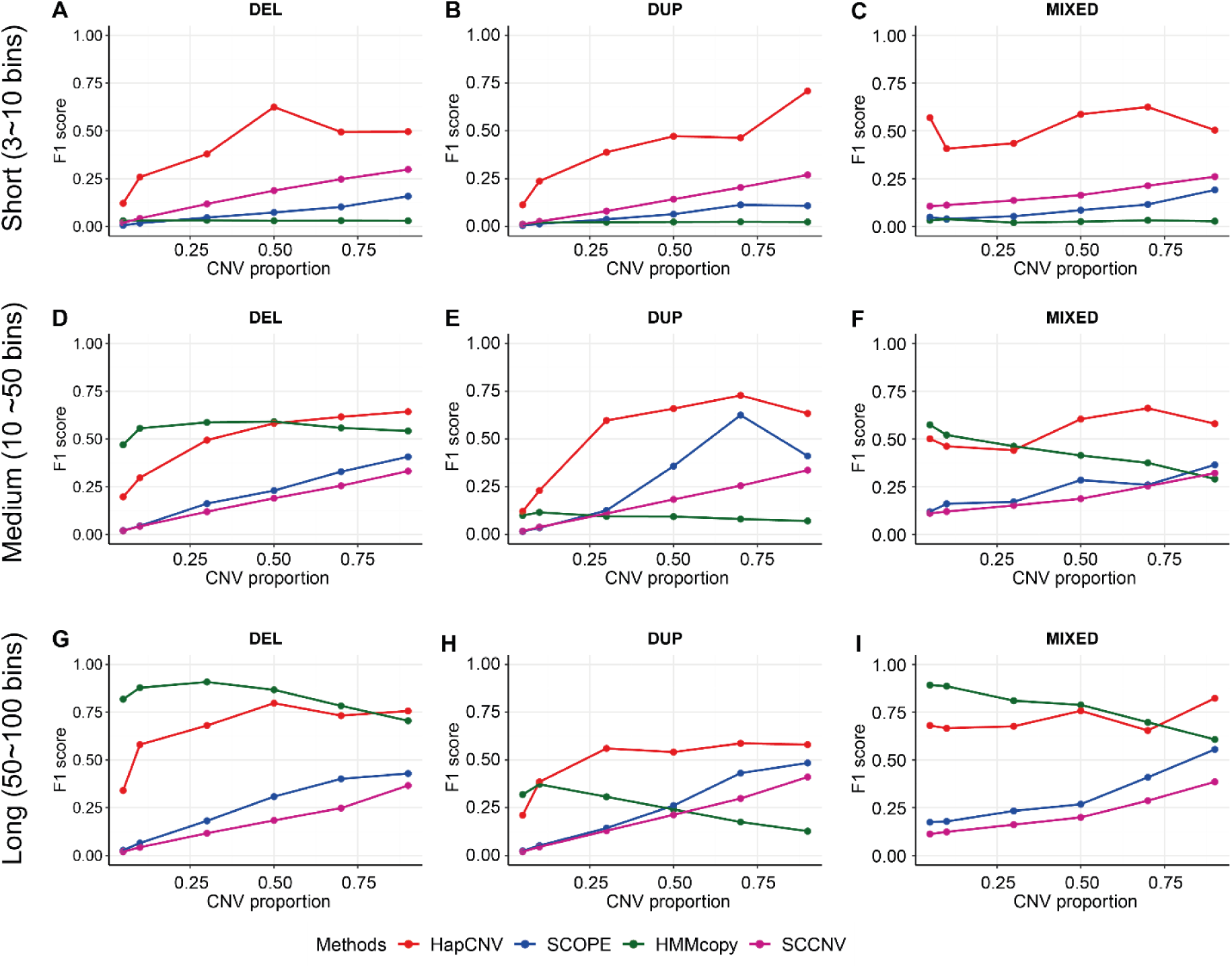
Performance evaluation of HapCNV using F1 score in comparison with existing methods for simulated data with amplification biases. For the simulations, we remained the locations of the genomic biomarkers prior to spiking in the CNV signals. Signals for 118 cells were simulated for three different CNV states: deletion of a single copy (DEL), duplication of a single copy (DUP), and mixture of two CNV states (MIXED) with varied length. The CNV proportion ranges from 5% to 90%.

### Analysis of low-cell sequencing data from the *P. falciparum* parasites

CNVs identified from the low-cell sequencing data of *P. falciparum* parasite were summarized in Table 1 and the updated posterior means of the identified CNVs were displayed in Supplementary Table 1. HapCNV, DNAcopy, and SCOPE displayed superior performance by identifying the known *pfmdr1* CNV in all samples, whereas HMMcopy and SCCNV only detected this CNV in a proportion of samples (19 and 11 of 27, respectively). For the known *pf332* CNV, HapCNV detected 18 out of 27 CNVs, outperforming HMMcopy and SCCNV, which identified 0 and 7 CNVs, respectively.

**Table 1.** The summary of identified CNVs from the low-input sequencing data from *P. falciparum*. Known: CNVs at known locations (*pfmdr1* CNV: chr5: 888001-970000; *pf332* CNV: chr11: 1950201-1962400). Common CNVs: CNV identified in > 10% of the cells. Rare CNVs: CNV identified in < 10% of the cells. There are 27 cells in total from the *Dd2* cell line, and all cells harbor the known CNVs. The table summarizes total number of detected CNVs and by CNV type (known, or common or rare).

| CNV type | HapCNV (%) | DNAcopy (%)<sup>a</sup> | SCOPE (%)<sup>a</sup> | HMMcopy (%) | SCCNV (%) |
| --- | --- | --- | --- | --- | --- |
| Known (*pfmdr1*) | 27 (2.6) | 27 (11.4) | 27 (0.8) | 19 (30.6) | 11 (0.6) |
| Known (*pf332*) | 18 (1.8) | - | - | 0 (0.0) | 7 (0.4) |
| Common | 917 (89.6) | 176 (74.3) | 2,563 (80.1) | 37 (59.7) | 1,620 (96.4) |
| Rare | 61 (6.0) | 34 (14.3) | 608 (19.1) | 6 (9.7) | 43 (2.6) |
| Total | 1,023(100.0) | 237 (100.0) | 3,198(100.0) | 62 (100.0) | 1,681(100.0) |

<sup>a</sup>Since DNAcopy and SCOPE need specified normal controls and the control cell line in our study had the *pf332* copy number variant, for fair comparison, we did not use them for *pf332* evaluation.

Consistent with the theory that the pseudo-reference construction in the normalization step will preferably retain the signals from common CNVs that are shared by reference and test samples, it was evident that HapCNV was advantageous in detecting common CNVs compared to other methods. For HapCNV, the detected common CNVs comprised a proportion of 89.6% of the total number of identified CNVs, which was the highest among the methods except SCCNV at 96.4% (Table 1). Consistently with the simulations, SCOPE and SCCNV detected a large number of total CNVs, possibly indicating their high susceptibility to amplification bias or weak CNV signals. For the rare variants, HapCNV also detected the second largest number just lesser than SCOPE, the latter was found to be sensitive and detected an overall excessive number of rare CNVs as well as common CNVs which was also shown in simulation studies.

To further elucidate the performance of HapCNV in detecting common CNVs from haploid genomes, we conducted a detailed assessment of the two known CNVs (*pfmdr1*, ∼82kb and *pf332*, ∼12kb, Table 2, Supplementary Fig. S6). By adaptively defining the baseline using pseudo-reference construction, HapCNV successfully detected *pfmdr1* CNVs in all samples. As we used the *FCR3* cell line (without the *pfmdr1* CNV) as the reference sample in the normalization step, as expected, methods relying on pre-specified reference samples including DNAcopy and SCOPE showed good performance in detecting *pfmdr1* CNVs. SCCNV, which employed across-cell normalization using *Dd2* samples, experienced CNV signal cancellation which led to a low recall rate of 0.259. Using within-cell information only, HMMcopy presented relatively low recall rate in detecting the *pfmdr1* CNV (Recall rate = 0.704; F1 score = 0.826). In detecting the *pf332* CNV (Table 2, Supplementary Fig. S6), HapCNV showed superior performance (F1 score=0.800), further validating the effectiveness of our method in preserving CNV signals without the need of pre-specified reference samples.

**Table 2.** Performance evaluation of methods in detecting known CNVs in *pfmdr1* and *pf332* with the *P. falciparum Dd2* parasite cell line. Precision rate was defined as TPs/(TPs + FPs); Recall rate as defined as TPs/(TPs + FNs). F1 score = 2*precision rate * recall rate/ (precision rate + recall rate). TP: true positives. FP: false positives. FN: false negatives. The true CNV status was through visualization of the signal intensities from the cells after normalization.

| CNV | Methods | Precision rate | Recall rate | F1 Score |
| --- | --- | --- | --- | --- |
| *pfmdr1* | HapCNV | 1.000 | 1.000 | 1.000 |
|  | DNAcopy | 1.000 | 1.000 | 1.000 |
|  | SCOPE | 1.000 | 1.000 | 1.000 |
|  | HMMcopy | 1.000 | 0.704 | 0.826 |
|  | SCCNV | 1.000 | 0.259 | 0.412 |
| *pf332* | HapCNV | 1.000 | 0.667 | 0.800 |
|  | HMMcopy | 0.000 | 0.000 | 0.000 |
|  | SCCNV | 1.000 | 0.111 | 0.200 |

## Discussion

The significant role of CNVs in explaining evolution and population diversity is gaining greater acknowledgment. Many studies have shown the connection between species evolution and the presence of rare or common CNVs [19–25, 55]. The availability of low-cell or single-cell DNA sequencing data makes it feasible to detect both rare and common CNVs at cellular resolution for various organisms, especially for those with haploid genomes. Although many statistical methods have been proposed for CNV detection, the reference samples used in normalization will bring bias to gene copy number evaluation. Additionally, some of these approaches may lose the ability to accurately detect CNVs within individual cells without leveraging information across cells [34].

In this paper, we developed the first CNV detection framework for haploid low-cell and scDNA-seq data. Different from traditional methods that select pre-defined whole cells as references, the unique bin-based pseudo-reference construction in the normalization step adaptively builds reference sequences for each cell, using information across cells from bins of normal copy number states thus reducing amplification biases and effectively reducing the amount of false discoveries of genetic variants [30, 35]. Our strategy utilizes a preliminary cell clustering method for unbiased baseline construction for normalization that will denoise the CNV signals after normalization and increases the overall accuracy and boost power in CNV detection, as reflected by our simulations and real data implications.

It was shown that HapCNV not only retains the power in detecting rare CNVs but also gains improved accuracy in detecting common variants. Comprehensive simulation studies demonstrated the program’s superior performance in detecting both rare and common variants, particularly for short CNVs (3∼10 bins). The analysis of haploid low-input data from *P. falciparum* also showed the enhanced power of HapCNV in detecting common variants. It is noteworthy that haploid and diploid scDNA-seq data share similar data structures that exhibit shallow and highly varied read count and amplification bias. Therefore, it is a natural extension to apply our method to diploid single-cell data with ploidy adjustment in the parameter settings.

By providing a natural and unbiased pseudo-reference construction for data normalization, our method has great implications for future computational development in relevant fields including copy number quantification in single-cell or spatial data. Nevertheless, HapCNV has some limitations. First, it relies on an initial cell clustering method (i.e., FLCNA), which adds additional computational time. Alternative methods such as traditional k-means or hierarchical clustering can be used but an additional step of shared CNV identification is needed. Second, due to the natural challenge of distinguishing amplification bias from weak CNV signals, noise sources cannot be completely removed from data by our newly developed normalization approach and some false positive calls will still be retained. This is not unique to HapCNV but also to all existing CNV detection approaches. Therefore, further efforts are still needed to effectively mitigate the effect of the amplification biases in future developments of copy number analysis approaches.

## CODE AVAILABILITY

The source code for HapCNV and examples of real data application can be accessed at https://github.com/FeifeiXiao-lab/HapCNV.

## Supplementary Files

**Supplementary Table S1.** The prior and posterior means in CNV detection for different copy number states from the *P. falciparum* genome. We generated the prior mean for each CNV state by clustering the segmental means detected by the circular binary segmentation (CBS) algorithm. Gaussian mixture model (GMM) was used to update the prior means using all segments and posterior means were obtained consequently. DEL: deletion of a single copy; Norm: normal state; Dup: duplication of a single copy.

| Copy number state | DEL | Norm | DUP |
| --- | --- | --- | --- |
| Prior mean | -1.798 | -0.052 | 2.489 |
| Posterior mean | -1.584 | 0.072 | 2.345 |

**Supplementary Fig. S1.**
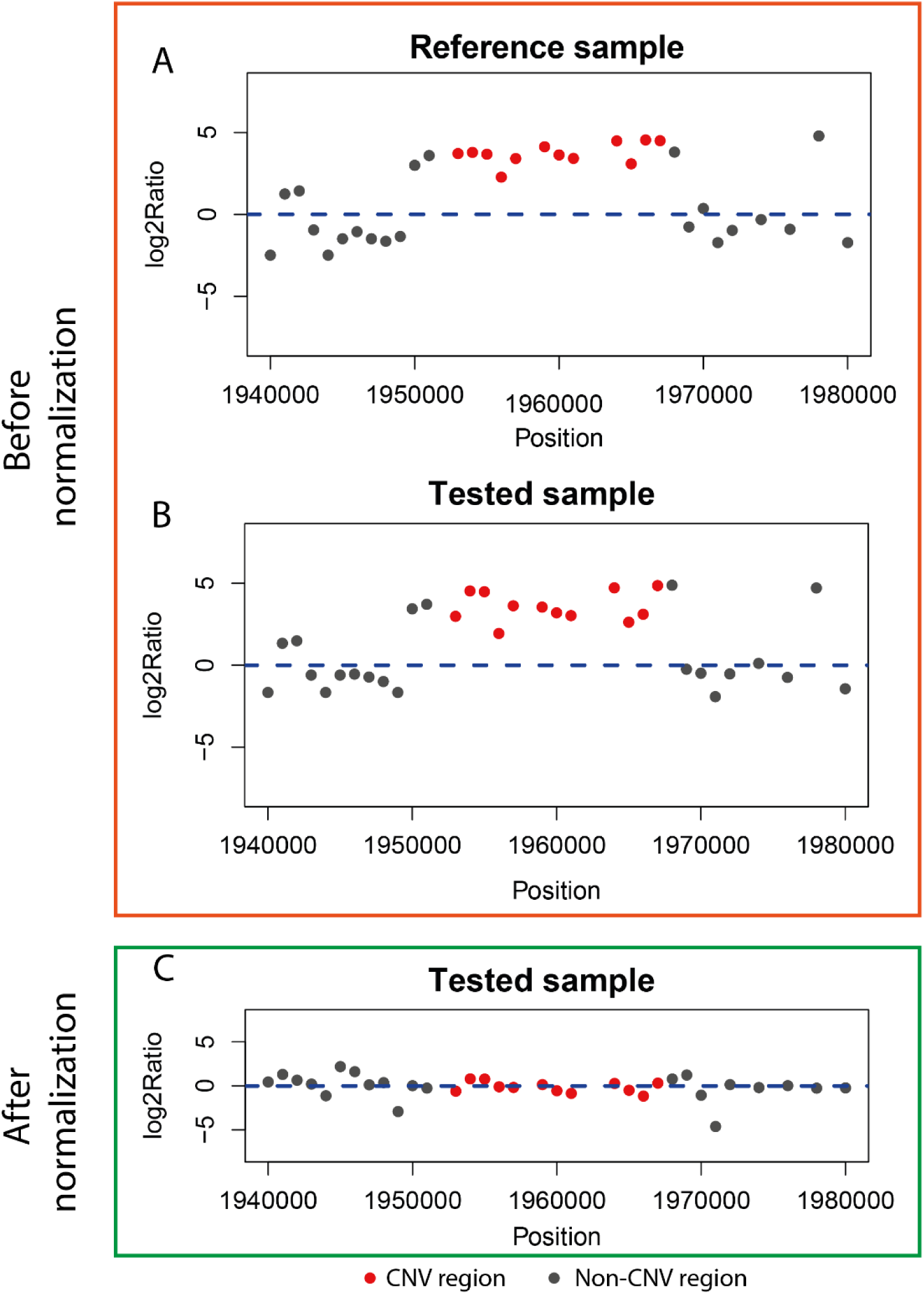
An example of using inappropriate reference that leads to signal cancellation. CNV signals within *pf332* gene are shown for a reference sample and a tested sample before and after normalization. Red dots denote the *pf332* CNV, while grey dots represent the neighboring non-CNV region. A, B: The reference sample is from the *FCR3* cell line. The tested sample is from *Dd2* cell line. Log2 ratios are the logarithmic transformation of the ratio between read counts and mean read count for each cell. C: Signals in tested sample are normalized by taking the logarithmic transformation of the ratio between read counts in tested sample and the median read counts from *FCR3* samples.

**Supplementary Fig. S2.**
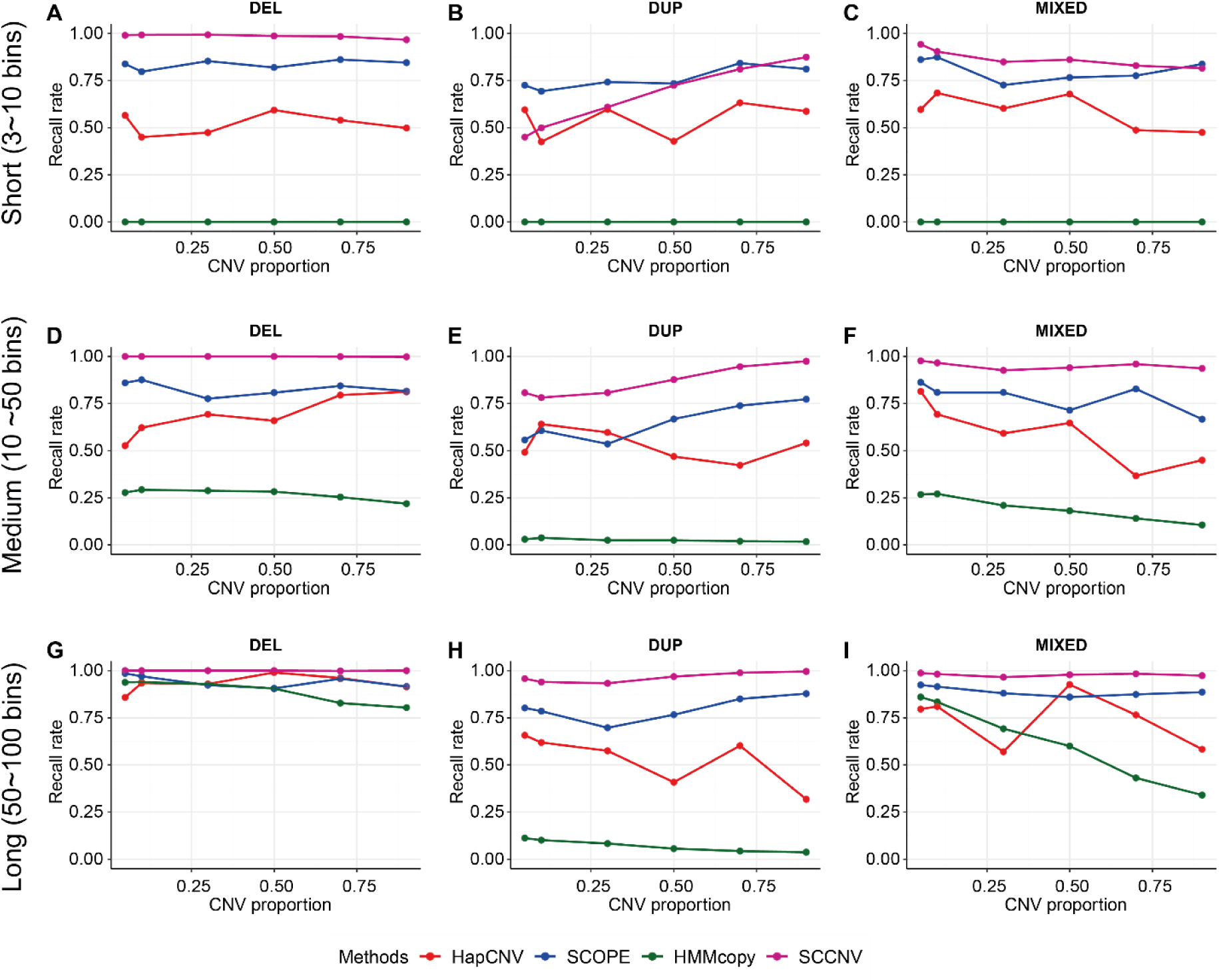
Performance evaluation of HapCNV using recall rate in comparison with existing methods for simulated data without amplification biases. For the simulations, we randomly shuffled the locations of the genomic biomarkers prior to spiking in the CNV signals to remove the effect that amplification biases may bring to the evaluation. Signals for 118 cells were simulated for three different CNV states: deletion of a single copy (DEL), duplication of a single copy (DUP), and mixture of two CNV states (MIXED) with varied length. The CNV proportion ranges from 5% to 90%.

**Supplementary Fig. S3.**
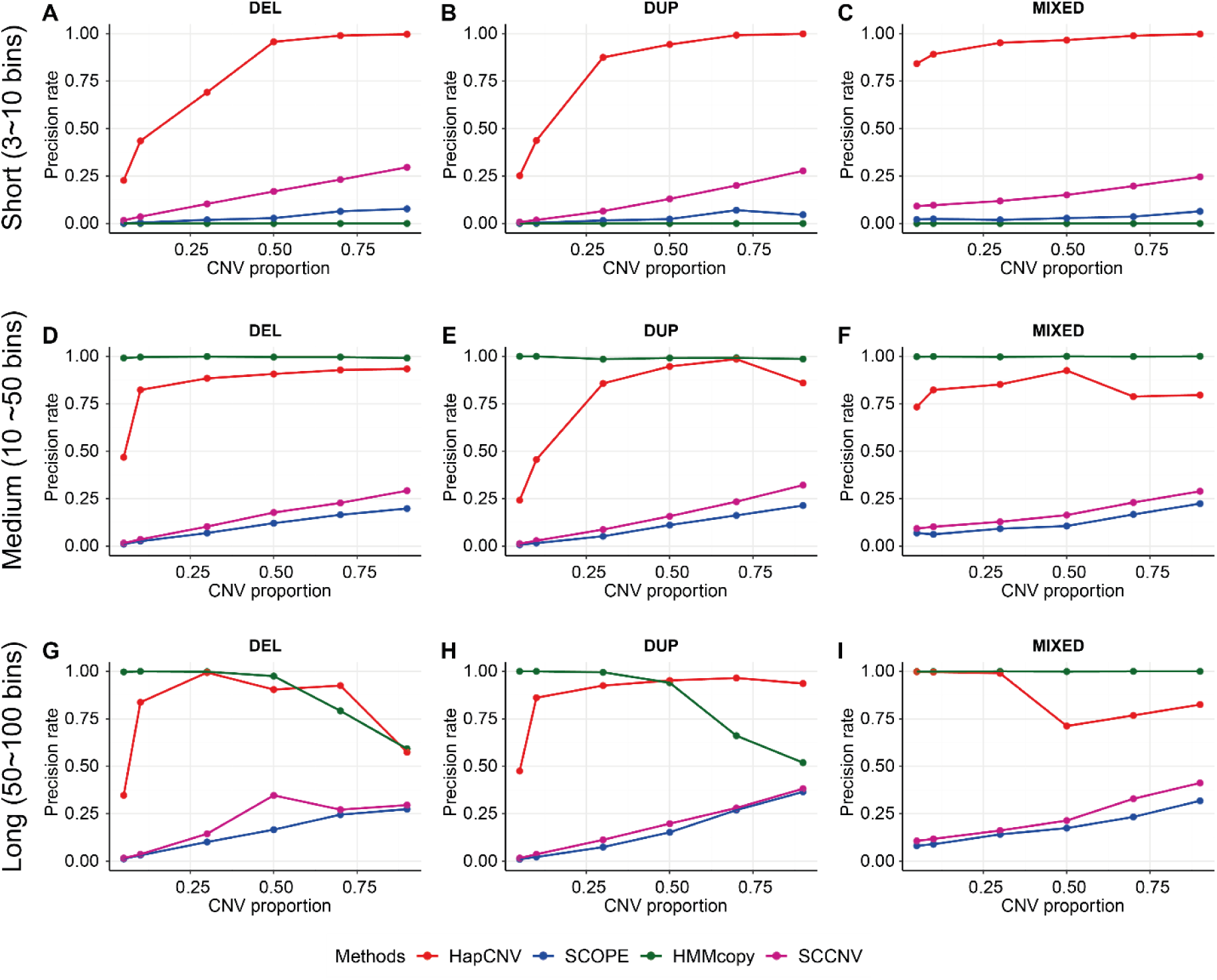
Performance evaluation of HapCNV using precision rate in comparison with existing methods for simulated data without amplification biases. For the simulations, we randomly shuffled the locations of the genomic biomarkers prior to spiking in the CNV signals to remove the effect that amplification biases may bring to the evaluation. Signals for 118 cells were simulated for three different CNV states: deletion of a single copy (DEL), duplication of a single copy (DUP), and mixture of two CNV states (MIXED) with varied length. The CNV proportion ranges from 5% to 90%.

**Supplementary Fig. S4.**
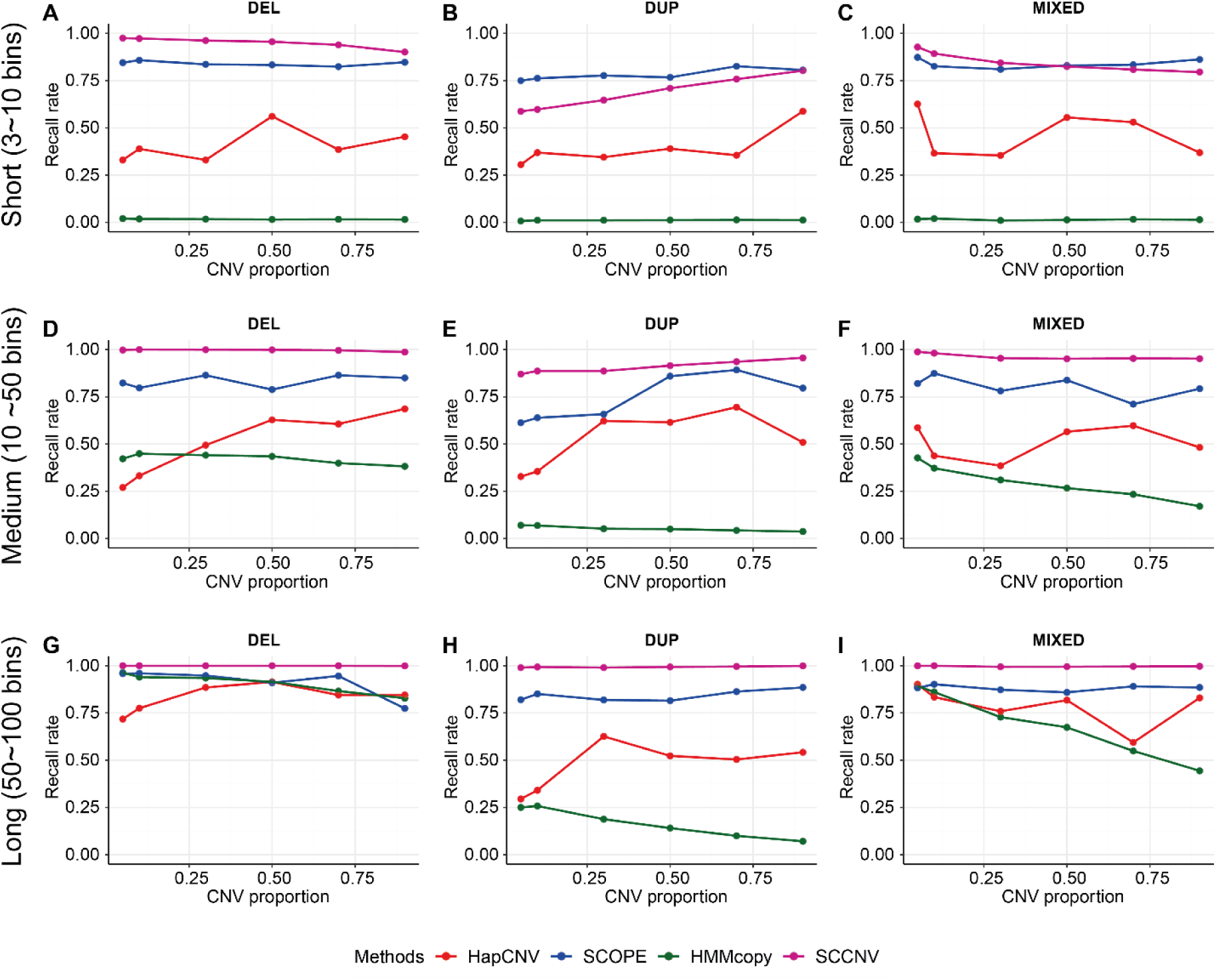
Performance evaluation of HapCNV using recall rate in comparison with existing methods for simulated data with amplification biases. For the simulations, the genomic structure of the background noise was mimicking the real dataset, so we assume the amplification biases remained in the data. Signals for 118 cells were simulated for three different CNV states: deletion of a single copy (DEL), duplication of a single copy (DUP), and mixture of two CNV states (MIXED) with varied length. The CNV proportion ranges from 5% to 90%.

**Supplementary Fig. S5.**
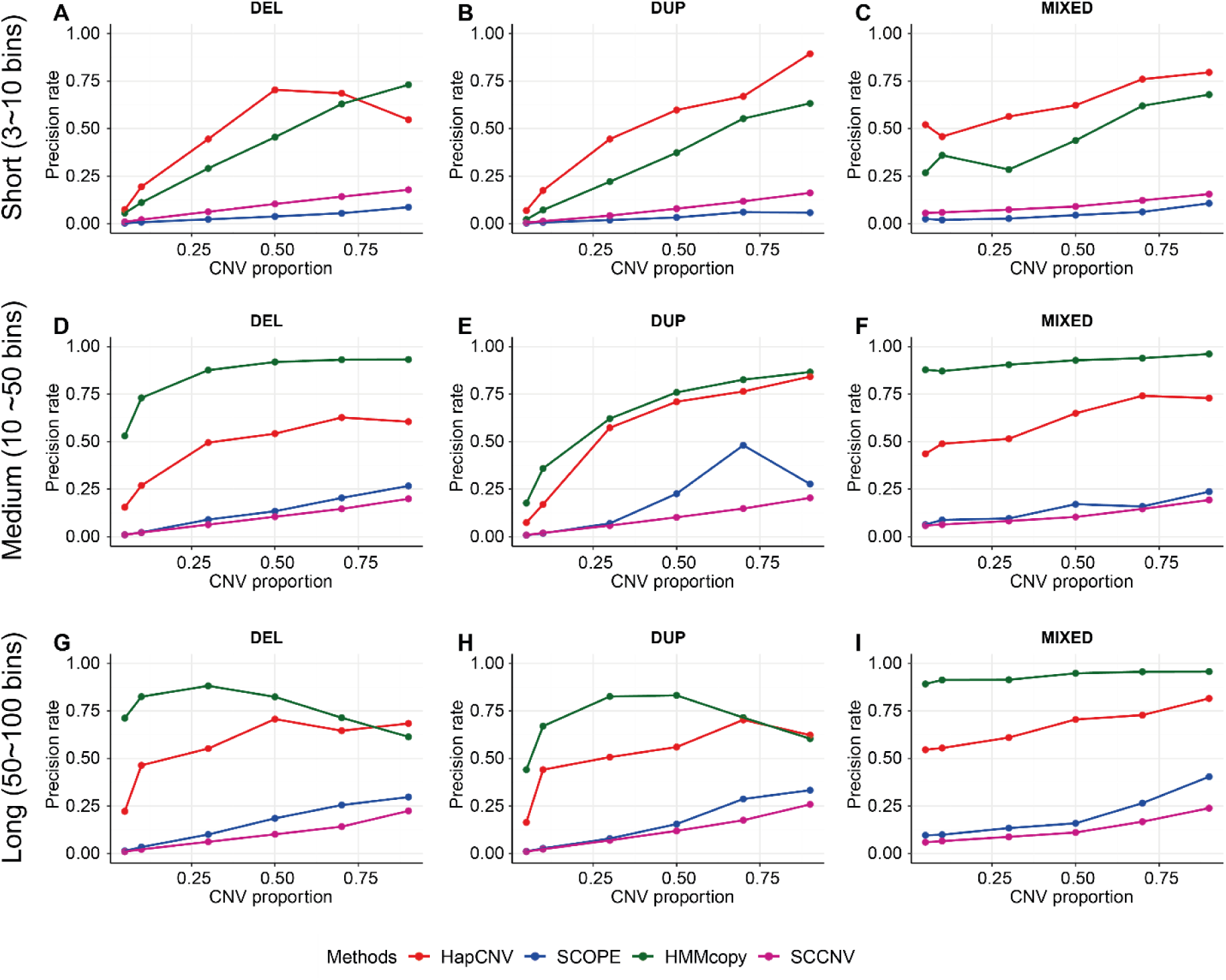
Performance evaluation of HapCNV using precision rate in comparison with existing methods for simulated data with amplification biases. For the simulations, the genomic structure of the background noise was mimicking the real dataset, so we assume the amplification biases remained in the data. Signals for 118 cells were simulated for three different CNV states: deletion of a single copy (DEL), duplication of a single copy (DUP), and mixture of two CNV states (MIXED) with varied length. The CNV proportion ranges from 5% to 90%.

**Supplementary Fig. S6.**
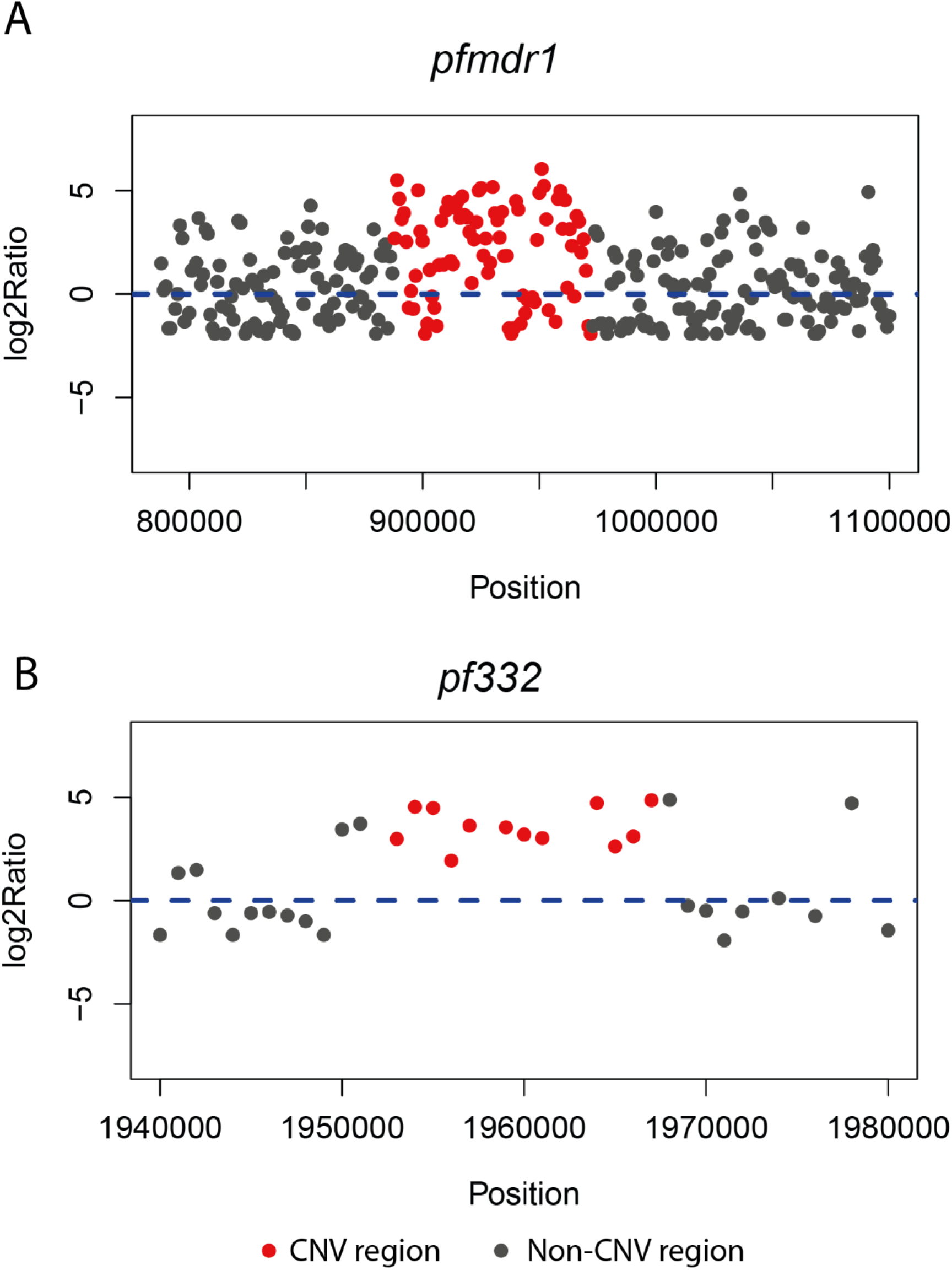
Signals for the *pfmdr1* CNV (Chr5: 888001-970000) and *pf332* CNV (Chr11: 1950201-1962400) from the *Dd2* parasite line. Red dots denote the CNV region, while grey dots represent the flanking regions. Log2Ratio: normalized read counts data by dividing the cell median.

## Reference

1. Tralamazza, S.M., et al., Copy number variation introduced by a massive mobile element facilitates global thermal adaptation in a fungal wheat pathogen. Nat Commun, 2024. 15(1): p. 5728.

2. Gschwind, A.R., et al., Diversity and regulatory impact of copy number variation in the primate Macaca fascicularis. BMC Genomics, 2017. 18(1): p. 144.

3. Guryev, V., et al., Distribution and functional impact of DNA copy number variation in the rat. Nat Genet, 2008. 40(5): p. 538–45.

4. Pereira, K.M.C., et al., Impact of C4, C4A and C4B gene copy number variation in the susceptibility, phenotype and progression of systemic lupus erythematosus. Adv Rheumatol, 2019. 59(1): p. 36.

5. Iskow, R.C., O. Gokcumen, and C. Lee, Exploring the role of copy number variants in human adaptation. Trends Genet, 2012. 28(6): p. 245–57.

6. Heo, Y., et al., Difference of copy number variation in blood of patients with lung cancer. Int J Biol Markers, 2021. 36(1): p. 3–9.

7. Yang, L., et al., Duplicated copy of CHRNA7 increases risk and worsens prognosis of COPD and lung cancer. Eur J Hum Genet, 2015. 23(8): p. 1019–24.

8. Krepischi, A.C., et al., Germline DNA copy number variation in familial and early-onset breast cancer. Breast Cancer Res, 2012. 14(1): p. R24.

9. Kumaran, M., et al., Germline copy number variations are associated with breast cancer risk and prognosis. Sci Rep, 2017. 7(1): p. 14621.

10. Hu, W., et al., Establishment of a novel CNV-related prognostic signature predicting prognosis in patients with breast cancer. J Ovarian Res, 2021. 14(1): p. 103.

11. Blount, Z.D., et al., Genomic analysis of a key innovation in an experimental Escherichia coli population. Nature, 2012. 489(7417): p. 513–8.

12. Kvitek, D.J. and G. Sherlock, Whole genome, whole population sequencing reveals that loss of signaling networks is the major adaptive strategy in a constant environment. PLoS Genet, 2013. 9(11): p. e1003972.

13. Gresham, D., et al., Adaptation to diverse nitrogen-limited environments by deletion or extrachromosomal element formation of the GAP1 locus. Proc Natl Acad Sci U S A, 2010. 107(43): p. 18551–6.

14. Baslan, T., et al., Genome-wide copy number analysis of single cells. Nat Protoc, 2012. 7(6): p. 1024–41.

15. Lang, G.I., et al., Pervasive genetic hitchhiking and clonal interference in forty evolving yeast populations. Nature, 2013. 500(7464): p. 571–4.

16. Lauer, S., et al., Single-cell copy number variant detection reveals the dynamics and diversity of adaptation. PLoS Biol, 2018. 16(12): p. e3000069.

17. Rich, S.M., et al., The origin of malignant malaria. Proc Natl Acad Sci U S A, 2009. 106(35): p. 14902–7.

18. Venkatesan, P., The 2023 WHO World malaria report. Lancet Microbe, 2024. 5(3): p. e214.

19. Liu, S., et al., Single-cell sequencing of the small and AT-skewed genome of malaria parasites. Genome Med, 2021. 13(1): p. 75.

20. Cheeseman, I.H., et al., Gene copy number variation throughout the Plasmodium falciparum genome. BMC Genomics, 2009. 10: p. 353.

21. Nair, S., et al., Adaptive copy number evolution in malaria parasites. PLoS Genet, 2008. 4(10): p. e1000243.

22. Ribacke, U., et al., Genome wide gene amplifications and deletions in Plasmodium falciparum. Mol Biochem Parasitol, 2007. 155(1): p. 33–44.

23. Veiga, M.I., et al., Globally prevalent PfMDR1 mutations modulate Plasmodium falciparum susceptibility to artemisinin-based combination therapies. Nat Commun, 2016. 7: p. 11553.

24. Price, R.N., et al., Mefloquine resistance in Plasmodium falciparum and increased pfmdr1 gene copy number. Lancet, 2004. 364(9432): p. 438–447.

25. Cheeseman, I.H., et al., Population Structure Shapes Copy Number Variation in Malaria Parasites. Mol Biol Evol, 2016. 33(3): p. 603–20.

26. Vande Zande, P., X. Zhou, and A. Selmecki, The Dynamic Fungal Genome: Polyploidy, Aneuploidy and Copy Number Variation in Response to Stress. Annu Rev Microbiol, 2023. 77: p. 341–361.

27. Liu, Z.L. and X. Huang, Copy number variants impact phenotype-genotype relationships for adaptation of industrial yeast Saccharomyces cerevisiae. Appl Microbiol Biotechnol, 2022. 106(19-20): p. 6611–6623.

28. Auwerx, C., et al., Rare copy-number variants as modulators of common disease susceptibility. Genome Medicine, 2024. 16(1): p. 5.

29. Pos, O., et al., DNA copy number variation: Main characteristics, evolutionary significance, and pathological aspects. Biomed J, 2021. 44(5): p. 548–559.

30. Wang, R., D.Y. Lin, and Y. Jiang, SCOPE: A Normalization and Copy-Number Estimation Method for Single-Cell DNA Sequencing. Cell Syst, 2020. 10(5): p. 445–452 e6.

31. Sabina, J. and J.H. Leamon, Bias in Whole Genome Amplification: Causes and Considerations. Methods Mol Biol, 2015. 1347: p. 15–41.

32. Brown, N., et al., Replication stress increases de novo CNVs across the malaria parasite genome. bioRxiv, 2024: p. 2024.12.19.629492.

33. Wang, X., H. Chen, and N.R. Zhang, DNA copy number profiling using single-cell sequencing. Brief Bioinform, 2018. 19(5): p. 731–736.

34. Lai D, H.G., Shah S, HMMcopy: Copy number prediction with correction for GC and mappability bias for HTS data. R package version 1.40.0. 2022.

35. Dong, X., et al., SCCNV: A Software Tool for Identifying Copy Number Variation From Single-Cell Whole-Genome Sequencing. Front Genet, 2020. 11: p. 505441.

36. Mills, R.E., et al., Mapping copy number variation by population-scale genome sequencing. Nature, 2011. 470(7332): p. 59–65.

37. Ravenhall, M., et al., An analysis of large structural variation in global Plasmodium falciparum isolates identifies a novel duplication of the chloroquine resistance associated gene. Sci Rep, 2019. 9(1): p. 8287.

38. Qin, F., et al., A statistical learning method for simultaneous copy number estimation and subclone clustering with single-cell sequencing data. Genome Res, 2024. 34(1): p. 85–93.

39. Magi, A., et al., EXCAVATOR: detecting copy number variants from whole-exome sequencing data. Genome Biol, 2013. 14(10): p. R120.

40. Xiao, F., et al., modSaRa: a computationally efficient R package for CNV identification. Bioinformatics, 2017. 33(15): p. 2384–2385.

41. Venkatraman, E.S. and A.B. Olshen, A faster circular binary segmentation algorithm for the analysis of array CGH data. Bioinformatics, 2007. 23(6): p. 657–63.

42. Zhang, Y., W. Liu, and J. Duan, On the core segmentation algorithms of copy number variation detection tools. Brief Bioinform, 2024. 25(2).

43. Miller, C.A., et al., ReadDepth: a parallel R package for detecting copy number alterations from short sequencing reads. PLoS One, 2011. 6(1): p. e16327.

44. Gusnanto, A., et al., Correcting for cancer genome size and tumour cell content enables better estimation of copy number alterations from next-generation sequence data. Bioinformatics, 2012. 28(1): p. 40–7.

45. Klambauer, G., et al., *cn.*MOPS: mixture of Poissons for discovering copy number variations in next-generation sequencing data with a low false discovery rate. Nucleic Acids Res, 2012. 40(9): p. e69.

46. Talevich, E., et al., CNVkit: Genome-Wide Copy Number Detection and Visualization from Targeted DNA Sequencing. PLoS Comput Biol, 2016. 12(4): p. e1004873.

47. Dharanipragada, P., S. Vogeti, and N. Parekh, iCopyDAV: Integrated platform for copy number variations-Detection, annotation and visualization. PLoS One, 2018. 13(4): p. e0195334.

48. Seshan VE, O.A., DNAcopy: DNA copy number data analysis. 2023. R package version 1.74.1.

49. Xiao, F., et al., An accurate and powerful method for copy number variation detection. Bioinformatics, 2019. 35(17): p. 2891–2898.

50. Scrucca L, F.C., Murphy TB, Raftery AE, *Model-Based Clustering, Classification, and Density Estimation Using mclust in R*. Chapman and Hall/CRC, 2023.

51. Layer, R.M., et al., LUMPY: a probabilistic framework for structural variant discovery. Genome Biology, 2014. 15(6): p. R84.

52. Garvin, T., et al., Interactive analysis and assessment of single-cell copy-number variations. Nat Methods, 2015. 12(11): p. 1058–60.

53. Gardner, M.J., et al., Genome sequence of the human malaria parasite Plasmodium falciparum. Nature, 2002. 419(6906): p. 498–511.

54. Kim, J.-H., et al., CNVRuler: a copy number variation-based case–control association analysis tool. Bioinformatics, 2012. 28(13): p. 1790–1792.

55. Zhao, W., et al., In vitro susceptibility profile of Plasmodium falciparum clinical isolates from Ghana to antimalarial drugs and polymorphisms in resistance markers. Front Cell Infect Microbiol, 2022. 12: p. 1015957.

